# AI-based identification of cardiac Purkinje fiber cells isolated from whole adult sheep hearts

**DOI:** 10.1101/2025.05.13.653917

**Authors:** Sebastien Chaigne, Thomas Hof, Kristell Mimoun, Ayoub El Ghebouli, Sabine Charron, Fabien Brette, Frank Choveau, Phillippe Pasdois, Catherine Rucker-Martin, David Benoist, Romain Guinamard, Michel Haissaguerre, Richard Walton, Olivier Bernus

## Abstract

**BACKGROUND:** Purkinje Fibers (PFs) are essential to the cardiac conduction system for synchronizing ventricular contractions. However, emerging evidence highlights their implication in the development of ventricular tachyarrhythmias. Nevertheless, isolating and studying the cellular mechanisms of PFs presents a significant challenge due to their intricate arborizing structure, heterogeneous cardiomyocytes (CMs) phenotype, and relatively small proportion within the ventricular mass, all of which hinder detailed functional investigations and comprehensive analysis of the conduction system network.

**OBJECTIVE:** To develop a new methodology for dissociation and classification of cell populations related to the ventricular conduction system from adult sheep. This workflow establishes, in part, a proof-of-concept deep learning-based classification strategy that leverages standard cellular imaging data.

**METHODS:** We developed a multi-tiered workflow to isolate and classify cardiac cell populations from adult sheep hearts. Coronary perfusion and enzymatic digestion were used to dissociate CMs from the left ventricular free wall (LVMs) and Purkinje-rich free-running false tendons (FTs). A three-pronged classification strategy was developed and implemented: (1) expert-guided visual phenotyping based on distinctive morphological traits; (2) rule-based morphometric quantification using automatic image analysis; and (3) deep learning-based classification with a retrained YOLOv9 model trained on augmented brightfield image datasets. This pipeline enabled accurate discrimination between LVM and FT-derived cells. Independent validation was performed using patch-clamp electrophysiology, T-tubule structure imaging with di-8-ANEPPS, and gene expression profiling (RT-qPCR) for Purkinje-specific biomarkers (Tbx5 and Cx40).

**RESULTS:** During the qualitative inspection, FT-dissociated cells had distinct morphological features, including an elongated or slender shape, finger-like projections, curves and tortuous shapes, and a new feature: the presence of spurs along the lateral membrane. Subsequently, a YOLOv9 model achieved an accuracy of 98% in distinguishing LVM and FT cells, based on the initial visual selection made by the operator. In addition, FT-cells exhibit a lower organization and density of T-Tubules compared to LVM. This classification was confirmed by the characterization of the typically longer action potential (AP) durations in FT cells. Finally, higher mRNA expression of the transcription factor Tbx5 and connexin40 (Cx40) was observed in FTs compared to left ventricular tissues.

**CONCLUSIONS:** We present a robust and scalable workflow for isolating and classifying cardiac Purkinje fiber cells from adult sheep, integrating manual phenotyping, rule-based morphometrics, and AI-driven deep learning. This multimodal approach enables high-accuracy identification of PF cells within heterogeneous tissue, confirmed through structural, molecular, and electrophysiological validation. Our findings overcome long-standing barriers in Purkinje fiber research and provide a powerful platform for advancing the study of ventricular conduction system biology and its role in arrhythmogenesis.

**GRAPHIC ABSTRACT:** 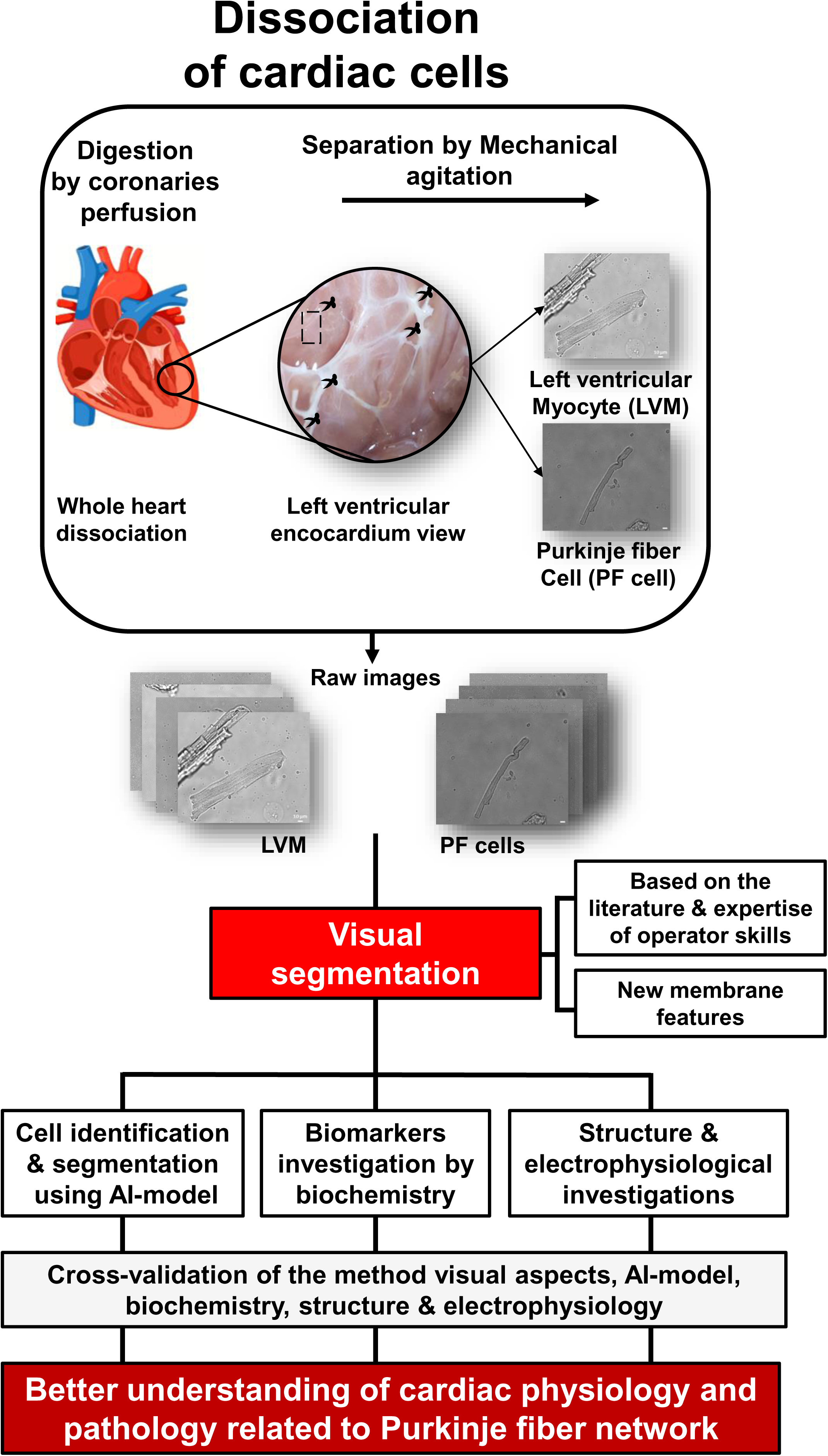

**WHAT IS KNOWN:**

- The PFs network constitutes a small part of the ventricular mass (<2%) but ensures spatio-temporal dynamic of ventricular activation.
- The PFs are known to have distinct electrophysiological and Ca^2+^ dynamic compared to surrounding myocardial tissue.
- Ventricular arrhythmias are the most common cause of sudden cardiac death (SCD), and recent evidence points to an essential contribution of PFs.
- However, little is known about the molecular mechanisms of PF-induced arrhythmias, partly because their isolation remains challenging.

**WHAT THE– STUDY ADDS:**

- A new dissociation technique combined with advanced AI methods to accurately dissociate and discriminate LVM and PF cells derived from free FT dissociation.
- This improves the classification system for distinguishing morphological LVM cells from PF cells in sheep.
- Our model opens up new perspectives in the automatic analysis of various cell parameters.

## INTRODUCTION

Purkinje fibres (PFs) are the terminal components of the cardiac conduction system, ensuring the fast and synchronized conduction of action potentials (APs) in right and left ventricles^1^. They have a unique embryological origin and specific electrical properties, including rapid AP conduction, prolonged AP duration, and specific management of Ca^2+^ handling^2,3,4,5,6,7^. Structurally, PFs exhibit a complex branching architecture that extends beneath the endocardial surface and into the ventricular transmural layer.^8^ Supporting a proportion of the PF network are free-running structures, known as False Tendons (FTs), which bridge trabeculae and papillary muscles in the ventricular cavity. Histological studies conducted across various species have previously revealed that FTs are composed of a combination of PF cells and an extensive extracellular collagen support that underlie the complexity of the specialized cardiac conduction system^9,10,11^. In addition, FTs can also contain varying levels of ventricular myocytes, nerve axons and blood vessels^10^. Compared to ventricular myocytes, PFs have fewer myofibrils, higher glycogen and mitochondrial content, and a less developed T-Tubule network, as has been observed in various species^5^. These structural and functional specializations reflect their unique role in the electrical conduction of the heart. It is increasingly-recognized that PFs play key roles in the development of arrhythmia leading to sudden cardiac death (SCD), responsible for ∼20-30% of all cardiac death^2,12^. Despite its importance in cardiac physiology and pathology, the conduction system remains poorly studied, and the cellular mechanisms underlying ventricular arrhythmias remain elusive, with no Purkinje-specific drug treatment available^2^. Unfortunately, due to several technical challenges^13,14^, there is currently no reliable method that guarantee high-yield isolation of quiescent and Ca^2+^-tolerant PF cells especially in large mammalian models. These challenges arise from the inherent heterogeneity of cellular phenotypes and the density of PF cells within the free FTs, which significantly limit the yield of viable PF cells after digestion. Furthermore, the delicate nature, complex organization, and intricate network of PF-cells make it extremely difficult to obtain an enriched population of quiescent PF cells for both structural and functional studies. This intricate structural arrangement, partially subsurface and deeply interwoven with CMs, makes PFs notoriously challenging to investigate. As such, there is a pressing need for improved methodologies that can overcome these limitations and enable robust, in-depth study of this elusive and highly specialized cardiac cell population.

To address this critical gap, we present in this report a newly-established dissociation and AI-method tailored for the isolation and discrimination of adult left ventricular myocytes (LVMs) and PF cells from adult sheep hearts. This innovative integrated strategy aims to improve the yield and viability of isolated PF cells derived from free FTs, which could lead to a better understanding of how these cells contribute to cardiac electrical activity and function. Furthermore, this AI-driven classification of FT-derived cells not only offers a transformative tool for cardiac cellular research but also sets a new standard for biomedical investigation, enhancing the accuracy, reproducibility, and scalability of future studies into cardiac pathologies.

## METHODS

### Bioethics

This study was carried out in accordance with the recommendations of the Directive 2010/63/EU of the European Parliament on the protection of animals used for scientific purposes and approved by the local ethical committee of Bordeaux CEEA50 (Ref: 21023).

### Experimental preparation

Hearts were obtained from 18 – 36 month-old adult female sheep. After pre-medication with ketamine (20 mg/kg) and acepromazine (Calmivet, 1 mL/50 kg), anesthesia was induced with intravenous injection of sodium pentobarbital (10 mg/kg) and maintained under isofluorane, 2%, in 100% O_2_. Heparin (2 mg/Kg) was injected intravenously to prevent blood coagulation. Sheep were euthanized by sodium pentobarbital (40 mL, from 50 mg/mL of stock) and the heart was rapidly excised. Finally, hearts were immersed in cold (4°C) Ca^2+^-free physiological saline solution (in mM): 140 NaCl, 5 KCl, 1 MgCl_2_, 10 HEPES, 10 glucose adjusted to pH 7.4 with NaOH.

### Cell isolation

Hearts were cannulated from both right and left coronary arteries (see Figure 1) and perfused (main infusion line 60 ml/min) for 10-15 minutes with Ca^2+^ free –physiological saline solution containing (in mM): 140 NaCl, 5 KCl, 1 MgCl_2_, 10 HEPES, 10 glucose adjusted to pH 7.4 with NaOH in order to flush and clean the heart cavities and coronary circulation of blood. Effective perfusion of circumflex and left anterior descending artery were critical for the experiment success. Then, hearts were perfused successively with an isolation solution containing (in mM): 130 NaCl, 5.4 KCl, 1.4 MgCl_2_, 0.4 NaH_2_PO_4_, 5 HEPES, 10 glucose, 10 creatine, 20 taurine (pH 7.6 with NaOH at 37°C) and supplemented with 0.1 mM EGTA for 15 minutes and collagenase A (0.239-0.252 U/mL, Roche) for a maximum of 30 minutes (pH 7.4 at 37°C). At the end of enzymatic dissociation, enzyme was cleared by perfusing the physiological saline solution containing 0.2 mM Ca^2+^. Atria were removed and ventricles were opened along the septum with caution. FTs were carefully (limiting FT and therefore PF cell stretching) dissected, avoiding ventricular myocardium at the insertion sites to limit contamination by CMs. Similarly, a 1 cm^3^ chunk of left ventricular free wall (LVFW) was also dissected for comparative cell isolation. Digested FTs and LVFW, were further minced with scissors, and mechanically shaken using a Pasteur glass pipette in isolation solution (0.2 mM Ca^2+^) to dissociate the cellular components (avoid bubbles during mechanical dissociation). Note that the experimental solutions were maintained under 100% O_2_ at 37°C during cell isolation steps. The corresponding cells were stored in Tyrode’s 1X solution with 0.2 mM Ca²⁺, ensuring optimal calcium levels that support cellular viability and function (LVFW vs. FTs from the right and left ventricles (see Figure S1)). Guided by brightfield microscopy, isolated cardiac cells were subject to further functional investigations and immunostaining.

**Figure 1:**
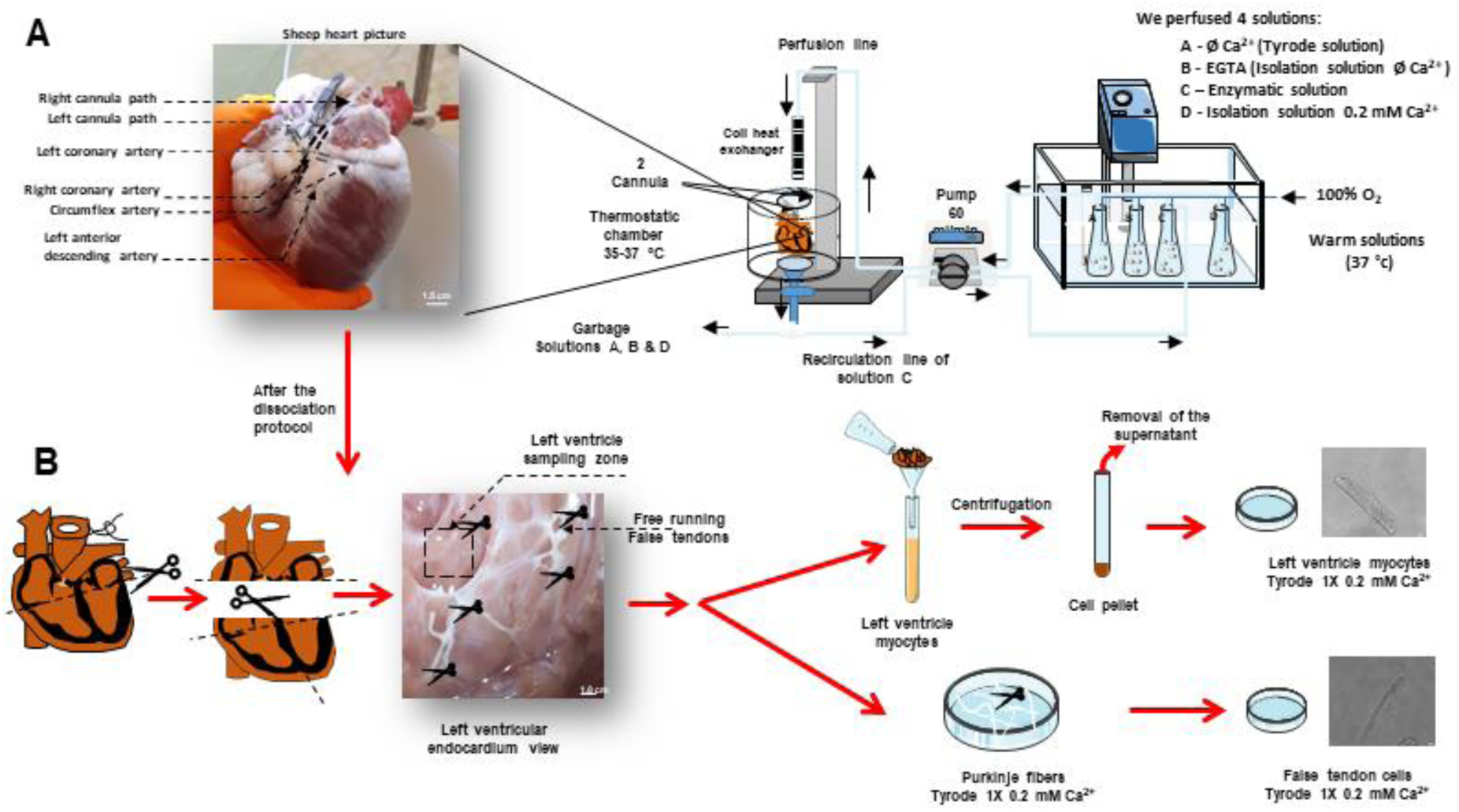
Cellular Dissociation Protocol. **(A)** Left: Representative view of the whole heart cannulated in the left and right coronary arteries. Right: schematic experimental dissociation setup. Perfusion solutions through the right and left coronaries were performed retrogradely to enable dissociation of both the left ventricular myocytes and Purkinje Fibers cells. **(B)** Left: ventricles were isolated from atria, and subsequently, the free running false tendons and the left ventricle myocytes were carefully removed, dissociated, and promptly immersed into a physiological solution with 0.2 mM Ca^2+^ solution.

### Cell imaging

For bright-field imaging, we used two distinct experimental setups. The first one used an inverted microscope (Nikon Eclipse Ti, with 40x magnification, Nikon instruments Inc, France) paired with a DS-Qi1Mc digital sight microscope camera (Nikon instruments Inc, France). The acquisition was realized by the NIS-basic research software (Nikon instruments Inc, France). The second one also includes an inverted microscope (IX83, with 60x magnification, Olympus, France) coupled with a CSU-W1 camera (Kakigama spinning disk, Andor Technology, UK) and MetaMorph software (Molecular Devices, USA). To record bright-field images, only rod-shaped myocytes that exhibited clear striations were used.

### Multifaceted morphological phentoyping

Brightfield images from the Nikon Eclipse Ti microscope of cells dissociated from both LVFW and FT regions (see Figure S2a) were subject to three distinct cell classification approaches. These included a manual morphological phenotyping, automatic morphometric analysis and AI-driven phenotype classification (Figure S2).

#### Qualitative visual classification

Expert-guidance confirmed the presence of whole striated rod-like cells which are consistent with cardiac muscle-derived cells within the image field of view in the brightfield images. Qualitative evaluation for previously described morphological features was applied to categorize cells based on the following characteristics: elongated/slender shapes^15,16,17,18^, finger-like projections^19^, curves^20,21,22^ and tortuous/tormented^15,22,23^ aspects of the plasma membrane. Qualitative inspection of the cell morphology provided manual validation of the inherent differences between cells dissociated from the FT and LVFW, and comparative qualitative morphological markers with existing literature describing PF cells and LVMs ^3, 4, 24, 25, 26, 27^.

#### Deep learning-based classification

In this study, we retrained the YOLOv9 model^28^ (yolov9e-seg) for the segmentation and classification of cells dissociated from the LVFW and FTs using an augmented dataset of 1998 images (the image bank owned by Liryc and linked to this study, which was initiated in 2018). This original dataset comprised 666 images manually validated to contain entire striated muscle cells within the image field of view. The image dataset was augmented with horizontal and vertical flips, rotations, brightness adjustments, exposure adjustments, and blur effects. The dataset was split into subgroups: 1846 images for training; 56 for validation, and 96 for testing. An epoch refers to one complete pass through the training dataset, while a batch represents a subset of the dataset processed in one iteration. In this context, the term “precision” indicates the relevance of results, and recall assesses how many relevant results are returned. High scores in both metrics indicate that the classifier is both accurate (high precision) and comprehensive in identifying positive results (high recall)^29^. Precision (P) is calculated as the number of true positives (𝑇𝑝) divided by the sum of true positives and false positives (𝐹𝑝): 𝑃 = 𝑇𝑝/(𝑇𝑝 + 𝐹𝑝)

Recall (R) is defined as the number of true positives divided by the sum of true positives and false negatives (𝐹𝑛): 𝑅 = 𝑇𝑝/(𝑇𝑝 + 𝐹𝑛)

The F1-score, the harmonic mean of precision and recall, is computed as: 𝐹1 − 𝑠𝑐𝑜𝑟𝑒 = 2𝑇𝑝/(2𝑇𝑝 + 𝐹𝑝 + 𝐹𝑛)

An example cell segmentation can be found in Figure S2b. A normalized confusion matrix was made for this cell classification model, which evaluated the performance of a classification model by comparing actual and predicted classifications.

#### Rule-based morphometric quantification

Cell segmentation outputs from the YOLOv9 model corresponding to cell regions of interest served for parallel automatic morphometric analysis. The segmented region was represented as a binary image of the same size as the original (Figure S2c; indicating the surface area in pixels). For length extraction, cells were often non-linear, requiring consideration for a curved longitudinal axis of the cell. Then, we applied the “skeletonize” function (Lee method), from the scikit-image library to the segmented region (Figure S2d). This function reduces binary objects to one-pixel-wide skeletons while preserving their topological properties^30^. Small skeletonized branches were eliminated to isolate the dominant longitudinal axis of the cell using the “FilFinder2D” class from the FilFinder package (Figure S2e), which is designed for analyzing filamentary structures in molecular clouds^31^. In order to optimize the length we applied smoothing of the skeleton using the “savgol_filter” function from the SciPy library (module scipy.signal) (Figure S2f)^32^. This function, provides smoothing for a one-dimensional data sequence while keeping its original shape. In our 2D image context, we applied it simultaneously to the X- and Y-axis coordinates of the skeleton in order to project the smoothed skeleton across the image plane. Finally, we extend the smoothed skeleton to the cell contour to establish the geodesic distance from each of the cell extremities (Figure S2g). Figure S2h displays the final median cell length curve (in pixels). To isolate a unique measurement of cell width, we divided the cell surface area by its length.

### Visualization of cell membranes and T-Tubules

To stain the cell membrane and observe T-Tubular organization and density, cells were incubated in the experimental chamber (tyrode with 0.2 mM Ca^2+^) with the lipophilic fluorescent indicator di-8-ANNEPPS (2 μM; Molecular Probes, Invitrogen, France) for 2 minutes and imaged using confocal laser scanning microscopy (Olympus France, Rungis, France) unit Yokogawa CSU-W1 (Japan) using 488-nm excitation light with detection at >505 nm. Quantitative analysis of T-Tubules organization was then assessed with “TTorg” plugin and specific script of ImageJ software (open source platform for biological image analysis (http://imagej.nih.gov/ij/))^33^.

### Tissue collection and RNA extraction

LVFW and FT samples were carefully dissected and immediately stored into liquid nitrogen. The corresponding biopsies were then stored at −80°C until RNA extraction. Total RNA was isolated from the tissues using the QIAzol reagent (QIAGEN), followed by purification and DNase treatment with the QIAGEN RNeasy Kit (QIAGEN). RNA quantification and purity were assessed by spectrophotometry (NanoDrop Technologies).

### Reverse Transcription quantitative PCR (RT-qPCR)

Sequences for primers were obtained from Ensembl Genome Browser. Primers were designed using Primer designing tool (NCBI) and synthesized at Sigma Aldrich / Merck. 200 ng of RNA was reversed transcribed using a cDNA Reverse Transcription kit (Ozyme - see below sequences) according to the manufacturer’s protocol. Quantitative PCR was performed in a 10 μL reaction volume (1 μL cDNA, 5μL of SYBR Green mix (Ozyme), a volume of 10 μM upstream and downstream primers respectively, and added double distilled water to 10 μL) on the BIO-RAD C100 Touch Thermal Cycler / CFX96 Real-time System. Real-time PCR conditions were as follows: 3 min at 95.0°C, 40 cycles of denaturation at 95°C for 5 s followed by 30 s annealing and elongation at 60°C. The primers used in this investigation were home-designed. Melting curves were obtained at the end of each run to confirm a single PCR product. All samples were run in triplicate. Non-template controls were introduced in each run to exclude contamination and nonspecific amplification. Expression levels of samples were normalized by using a normalization factor calculated by CFX Manager software (BIO-RAD). This normalization factor was calculated based on RT-qPCR results for two selected reference genes, GAPDH and GUSB. This allowed quantification of the target gene in one sample relative to that in another (the calibrator) using the “2 −ΔΔCtmethod” of calculating fold changes in gene expression.

Primer sequences of our biomarkers for real-time PCR studies: Tbx5 and Cx40 respectively (F: Forward primer and R: Reverse primer):

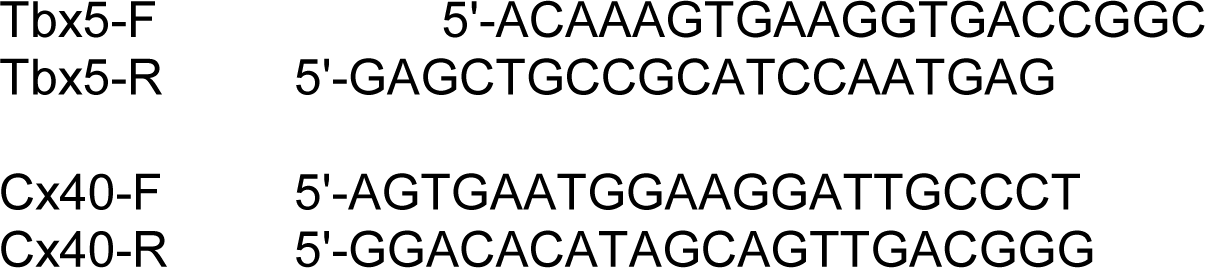

### Patch-clamp experiments and solutions

Isolated cells from the LVFW or FTs were placed in a petri-dish on an inverted microscope (Eclipse Ti-U, Nikon Instruments Inc, France). Patch-clamp experiments were all conducted at physiological temperature (∼37°C) and using whole cell configuration. APs were triggered at 1 Hz by a current square pulse (from 300 to 1,200 pA for 2 ms). Resting membrane potential (RMP), action potential amplitude (APA), maximal upstroke velocity (Vmax), and APDs at 30, 50, 70, and 90% of repolarization (APD30, APD50, APD70, and APD90) were monitored. Data acquisition and analysis were performed with Patchmaster and Fit Master (HEKA Elektronik, Lambrecht, Germany). Pipettes were pulled from borosilicate capillary glasses (W150F, WPI, UK) with a DMZ universal puller (Zeitz Instruments, Germany). The resistance of the filled pipettes was 1–2 MΩ. For AP recording, pipette solution contained (in mM): 110 K-aspartate, 25 KCl, 5 NaCl, 3 Mg-ATP, 10 HEPES, 0.002 cAMP, 10 PhosphoCreatine-K_2_, 0.01 EGTA. External solution contained (in mM): 140 NaCl, 5 KCl, 1 MgCl_2_, 10 HEPES, 10 Glucose and 1 CaCl_2_ (pH 7.4 NaOH).

### Statistics

Data are presented as mean ± Standard Error of the Mean (S.E.M). GraphPad Prism (GraphPad Software (version 7.04, (GraphPad Software, Boston, MA, USA)) was used for the statistical analysis and graph creation. Datasets were tested for normality using simultaneously d’Agostino-Pearson test and student’s t-test or one-way ANOVA. Differences with a p value of <0.05 were considered statistically significant.

## RESULTS

### Anatomy observation

The ventricular conduction system in sheep contains sophisticated arrangement of PFs, distributed between proximal free-running (false tendon [FT]) structures and an intricate distal network running endocardially and transmurally. We observed a more extensive FT network in the LV compared to the RV in sheep (Figures 1 & S1), which is consistent with previous reports from different mammalians species^26,34^. This more pronounced development of the PF network in the LV underscores its enhanced role in the overall cardiac rhythm and function ^35,36^.

### Morphology and dimensions of sheep left ventricular myocytes and false tendon cells

Whole hearts from healthy sheep were perfused with an enzymatic solution to digest cardiac tissue (Figures 1A). Due to their exposure in the LV cavity and ease of identification, central portions of FTs were dissected and treated for cell isolation using mechanical agitation. Cardiomyocytes were isolated in parallel from dissected tissue sections of approximately 1 cm^3^ from the LVFW for equivalent agitation practices (Figure 1B). Cells isolated from both FTs and LVFW were tolerant to Ca^2+^ and maintained viability for a duration approximating 8-10h post-dissociation. Cells dissociated from both regions were suitable for conducting morphological, structural, biochemical and functional studies. As cells isolated from FTs are not representative of the entire PF network, from this point onwards, we refer to cells isolated from FTs as FT-dissociated cells.

### Manual morphological phenotyping

We qualitatively identified distinct morphological phenotypes that differentiates LVM and FT cells, highlighting notable changes in membrane-specific aspects and cell shape (see Figures 2A & 2B). Compared to LVMs, FT cells showed a slender profile, sometimes with an elongated shape or the presence of finger-like projections at the end of cells, tortuous or tormented longitudinal axes, pronounced curvatures as well as spurs, on their plasma membranes. Note that similar striations to those in the LVM have been observed previously in sheep FT cells.

**Figure 2:**
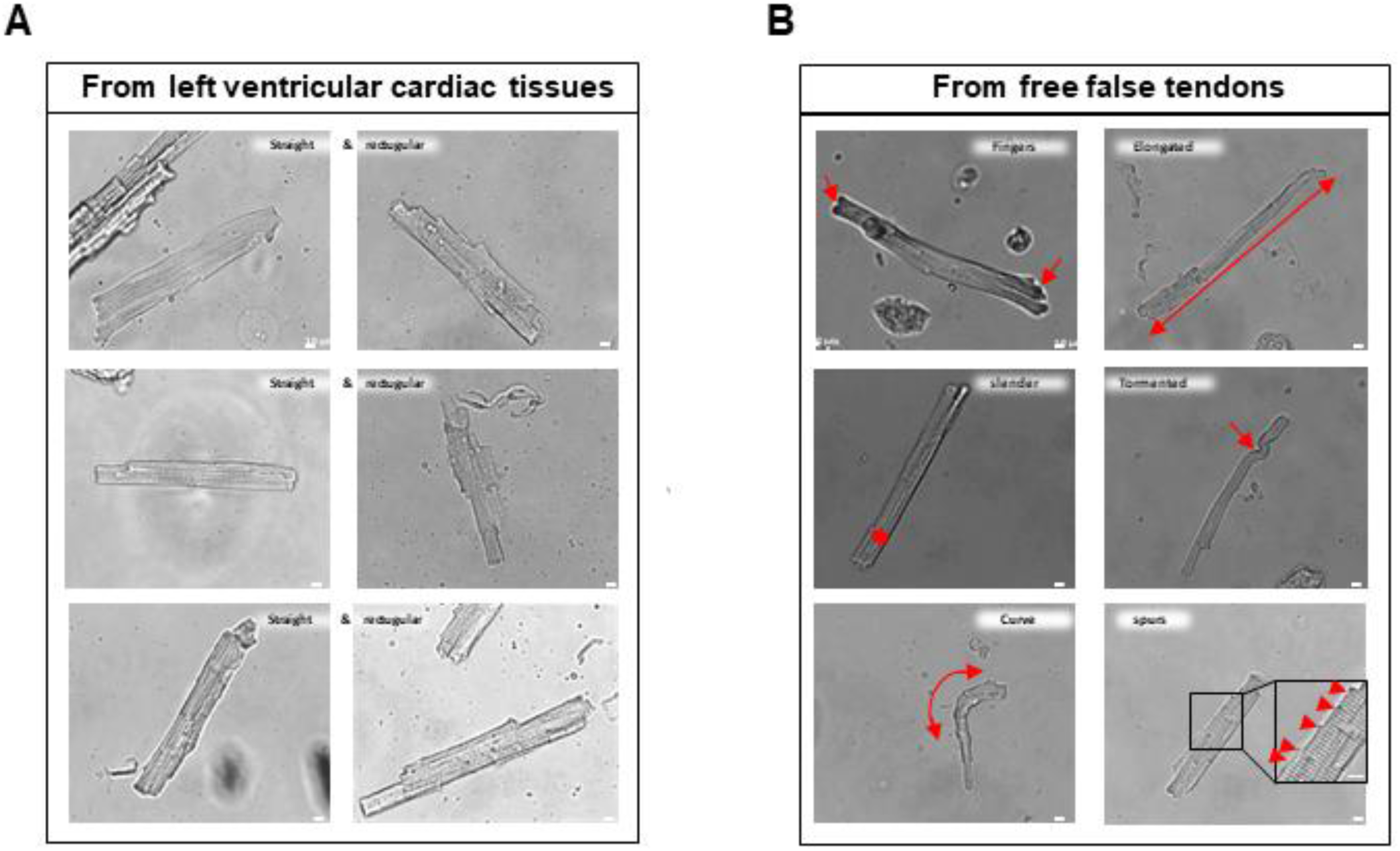
Morphological comparison between left ventricular cardiomyocytes and False tendon cells from adult sheep heart. **(A)** Typical shape of left ventricular myocytes (LVM - mainly straight and rectangular). **(B)** False tendon cells (FT cells) exhibit specific morphological aspects such as an elongated shape, slender size, spurs, tortuous appearance, the presence of finger-like projections at the intercalated discs level, and curvatures.

### Cardiac Cell Phenotype Classification Using Deep Learning

Our retrained YOLOv9 model demonstrated high performance in classifying cardiac cell types, achieving a precision of 99.1%, recall of 98.8%, and an F1-score of 98.95% on the validation dataset after 454 training epochs (Figure 3A). The normalized confusion matrix confirmed the model’s ability to accurately discriminate between LVM and FT cell populations (Figure 3B). This high classification accuracy validates the utility of deep learning in distinguishing morphologically similar cell types and provides strong support for the automated phenotypic separation of experimental groups, thereby reducing human bias and increasing throughput in cardiac cellular studies.

**Figure 3:**
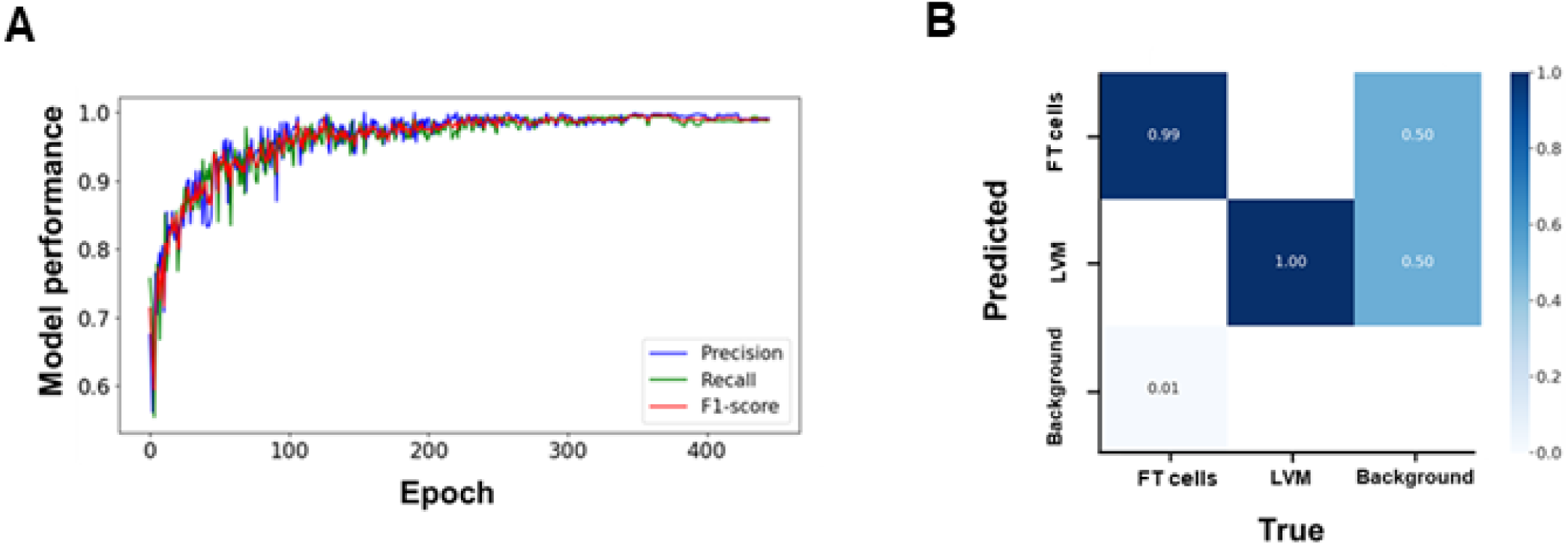
Cardiac Cell Phenotype Classification Using Deep Learning (YOLOv9 model training): (**A**) Precision, recall, and F1-score progression throughout the training process; (**B**) Normalized confusion matrix.

### Rule-based morphometric quantification

The integrated morphological classification framework implemented parallel rule-based analysis of cell morphology using a cell segmentation (binary image) output from the retrained YOLOv9 model. Figure 4A & Table S1 show comparative assessments of LVM and cardiac FT cell morphology based on selected features. FT cells have significantly thinner morphology and a reduced surface area (percentage differences of mean values were 38.9% and 28 %, respectively). However, cell length was statistically unchanged between cells from each region.

**Figure 4:**
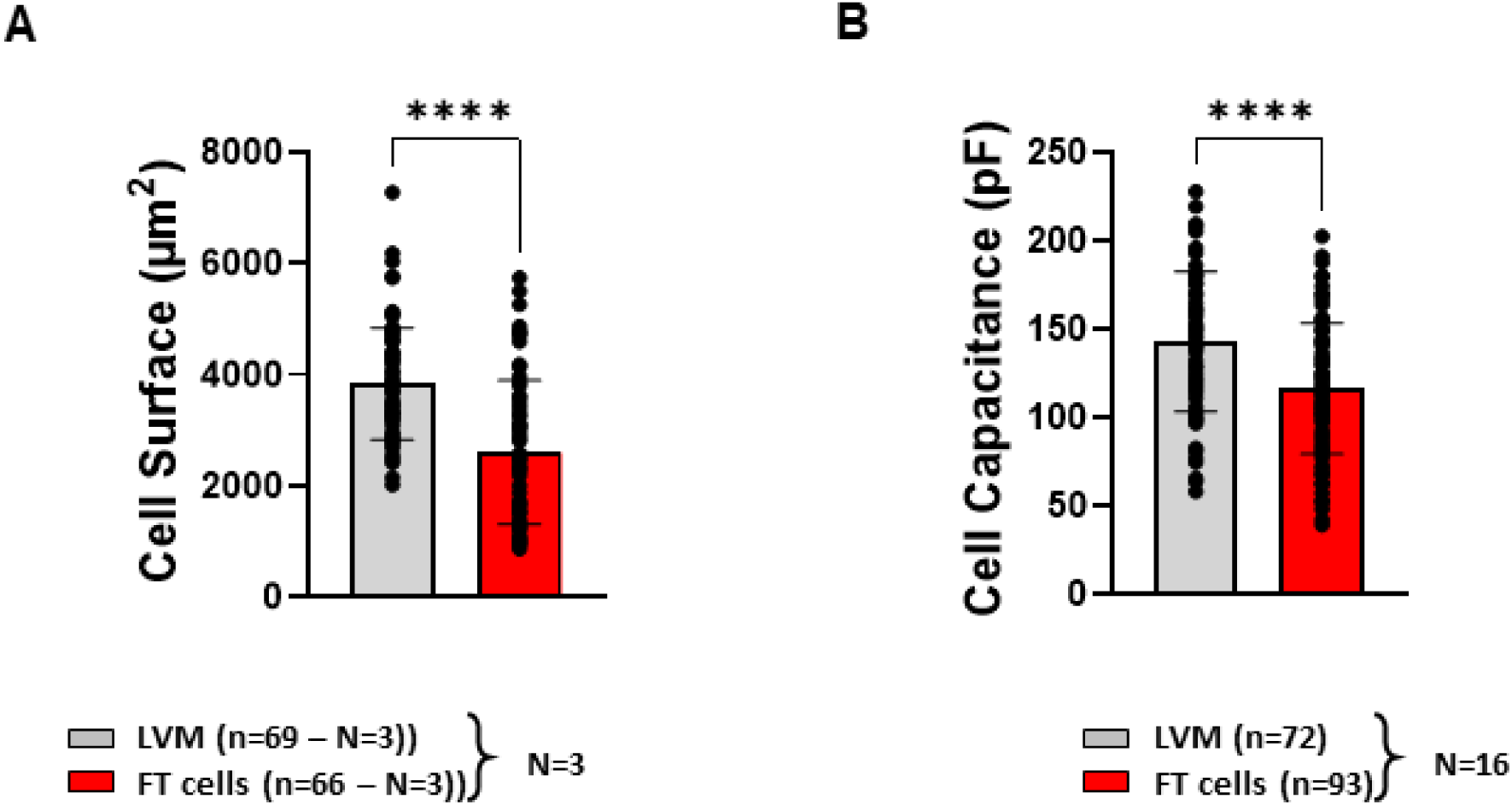
Cell Surface and Capacitance. (**A**) Average surface area of left ventricular myocytes (LVM) and false tendon (FT) cells under control conditions, evaluated by automated proceedings. (**B**) Average capacitance of LVM and FT cells under control conditions, evaluated by whole-cell patch-clamp recordings. Data are represented as mean ± standard error of the mean with individual values for each cell. N indicates number of animals; n, number of cells. Unpaired t-test, ****p < 0.0001.

Consistent with semi-automated morphometric analysis, cellular patch-clamp recordings provided independent and comparative measurements of cell membrane surface area by assessing cellular electrical capacitance. In agreement with morphological measurements, a 21 % lower capacitance was found in FT cells compared to LVM (Figure 4B). Note that both cellular groups maintained their initial appearance for 8-10 hours in room temperature post dissociation.

### Specific biomarkers of false tendon cells

It is known that the maturation process and function of the PF during embryonic stages involves several transcription factors, including Nkx2-5 and Tbx-5^3,4^, both of which have specific expression patterns in these specialized cells^37^. In addition, Connexin 40 (Cx40), a critical gap junctional protein in the PF network responsible for rapid impulse propagation between cells ^38^. We performed q-PCR studies and confirmed the presence of higher Tbx5 and Cx40 mRNA expression in FT cells compared to LVM, which highlights this differential expression pattern (Figure 5). These findings align with previous studies, showing that Tbx-5 and Cx40 are specific biomarkers of FT cells^3,4,5,26,37,39^.

**Figure 5:**
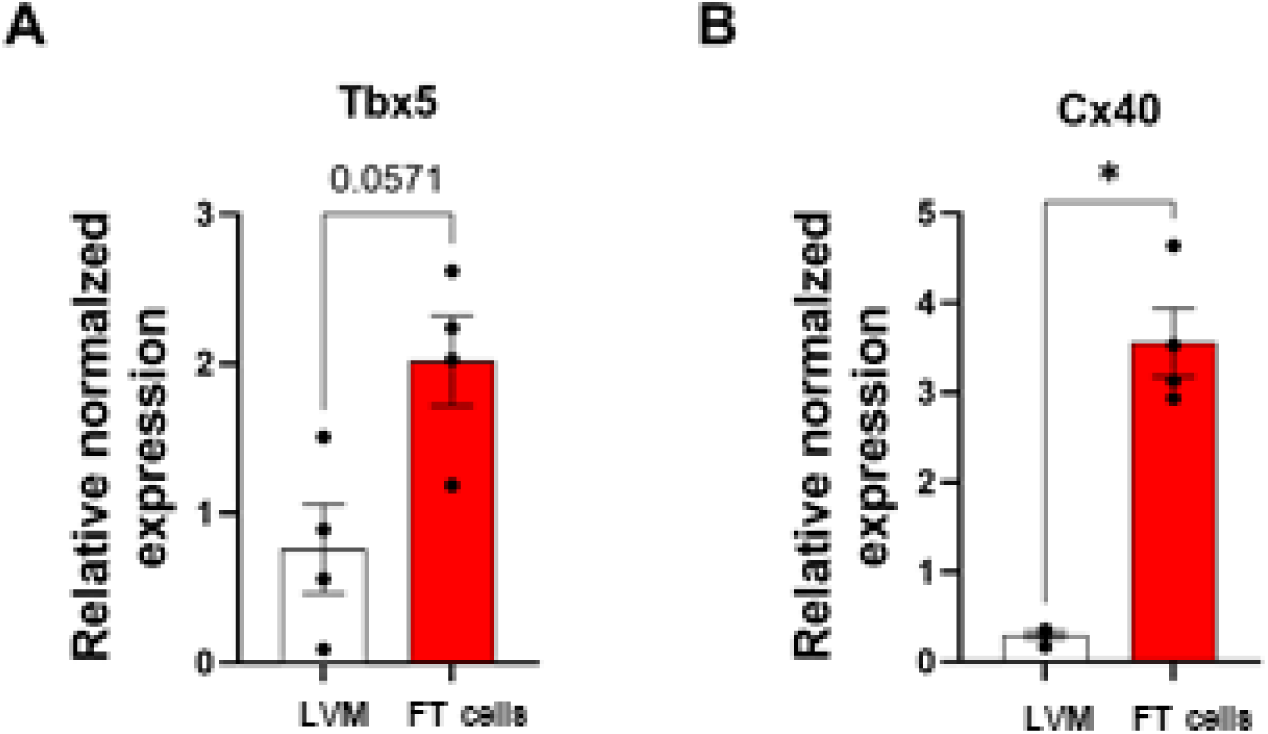
Expression of T-box transcription factor 5 (Tbx5) and Connexin 40 (Cx40) genes in the left ventricle myocytes and false tendon cells of adult sheep. Bar graphs illustrating quantitative results obtained through q-PCR to evaluate the expression of Tbx5 (**A**) and Cx40 (**B**) genes in left ventricular myocytes (LVM) and false tendon (FT) cells of adult sheep. The results suggest both a higherTbx5 and Cx40 mRNA expression in FT cells compared to LVM.

### T-Tubule Network organisation

The organization and heterogeneity of the T-tubule network was investigated by labeling the plasma membrane and T-Tubular network with di-8-ANNEPPS followed by confocal microscopy imaging to assess structural differences between LVM and FT cells. Firstly, the Fourier transformation decomposes excitation-contraction T-tubule images into their frequency components, thereby facilitating the quantification of repetitive and regular structures. In instances where mathematical analysis was not possible, we reported the percentage of cells that permitted effective quantification of level of T-tubules organization (LVM 87.3% vs. FT cells 70.8% of success). The lack of results was attributed to insufficient or no organization (see Figure 6-left). FT cells displayed a significant lower T-Tubular organization (see Figure 6) in comparison to LVM. The observed differences suggest non-uniform structural organization of the T-Tubule system throughout the cardiac muscle, which can have implications on the efficiency of excitation-contraction coupling and overall cardiac function. Then, we used ImageJ software to simultaneously extract the skeleton of each image and applied a script that defines the degree of luminosity within each cell for T-Tubule density quantification. The corresponding results suggest that T-Tubule density is significantly lower in FT cells compared to LVM. In this context, the more organized and denser T-Tubule network in LVM explains the better Ca^2+^ regulation and stronger contractile responses compared to the less organized, less dense network in FT cells (Figure 6, right). Finally, we observed no significant differences in the T-tubule distances between the two cell types.

**Figure 6:**
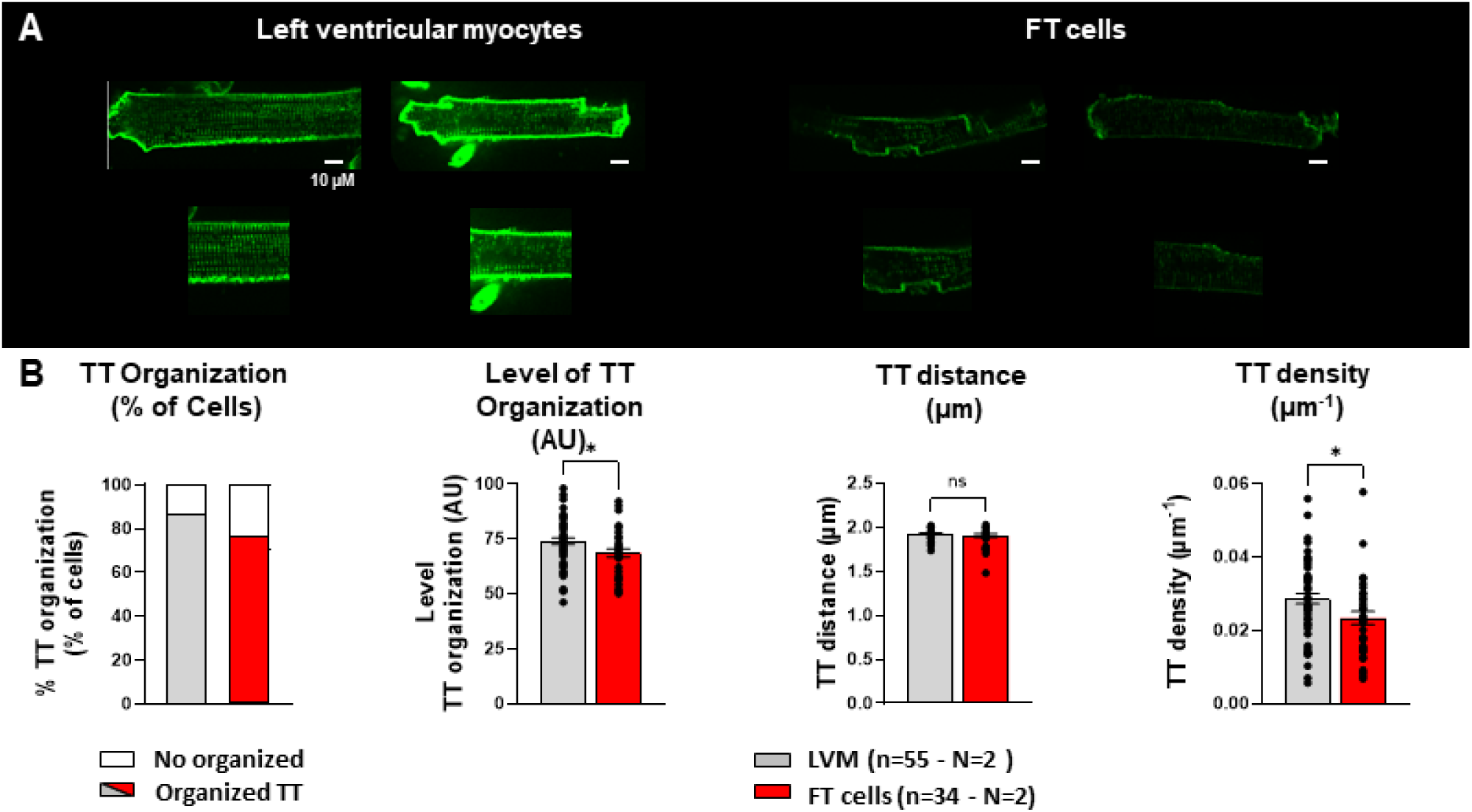
Representative images of isolated left ventricular myocytes and False tendon cells stained with di-8 ANNEPS and visualized under confocal microscopy. (**A**) T-Tubule characteristics for both left ventricular myocytes (LVM) and false tendon (FT) cells in adult sheeps. 10% magnification for each image are also shown below. (**B**) Percentage of cells showing an organized T-Tubule network (grey and red bars) or a non-organized T-Tubule network (white bars). Mean ± SEM T-Tubule regularity in organized cells and associated density. Data for T-Tubule organization and density analysis are realized under ImageJ software. N indicates number of animals; n, number of cells. Unpaired t-test, * p < 0.05.

### Cardiac cell electrophysiology

In our study, we compared the AP profiles between LVM and FT cells (Figure 7 & table S2) to characterize the electrophysiological specificities, unique to each of these cardiac cell types. This analysis allowed us to differentiate both cell types based on their electrophysiological properties. Please note that cells isolated from the LVM required a stabilization period (∼4hours) before functional exploration. Indeed, the methodology was tailored to selectively extract cardiac FT cells according to their microenvironment and ensure accessibility to LVM despite a higher degree of digestion. Compared to LVM, FT cells APs exhibit significant prolonged AP duration (APD_90_: 336.8± 24.7 vs 435.7 ± 26.00 ms* (Figures 7A & B). No significant differences were observed in the maximal upstroke velocity (dV/dt max: 212.8 ± 31.7 vs 236.9 ± 21.8 V/s (Figure 7C & table S2), amplitude (APA: 106.5 ± 3.1 vs 112.30 ± 2.4 mV (Figure 7C and table S2) or in other electrophysiological properties such as the phase 1 of repolarization profile, resting membrane potential and plateau voltage amplitude linked to the phase 2 (Figures 7A, S3, S4 and table S2). It is noteworthy, however, that mean values of dV/dt max and APA tended to be higher in FT cells, despite the lack of statistical significance. Finally, the AI-assisted post-processing of cardiac cell images (LVM vs. FT cells) from the functional investigation revealed also distinct phenotypes within our two experimental groups (see Figure S5). This confirms the efficiency of our cell identification and classification method.

**Figure 7:**
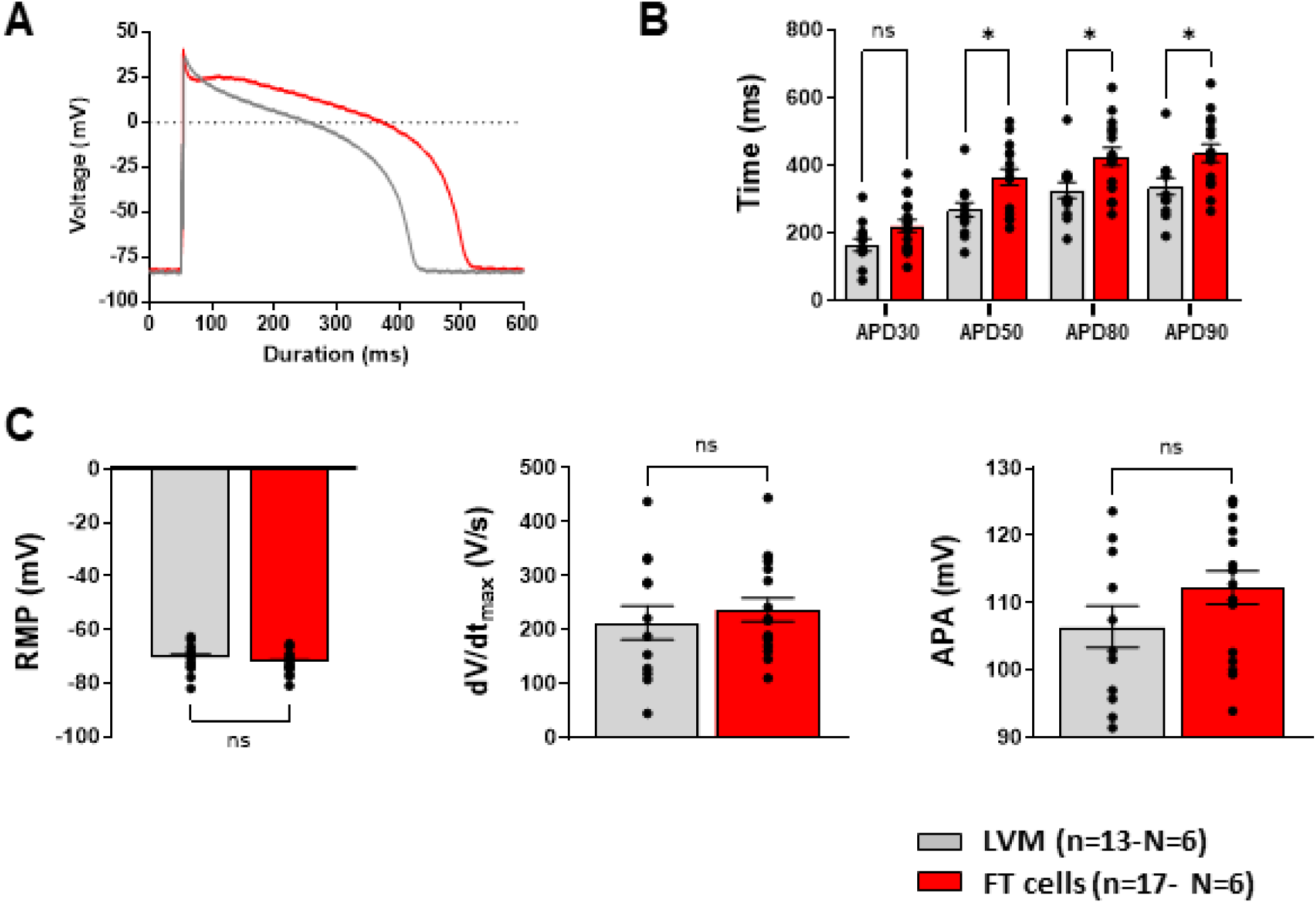
Comparison of action potential parameters from left ventricular myocytes (grey) and False tendon cells (red) in adult sheep. (**A**) Representative action potentials from isolated left ventricular myocytes (LVM) and false tendon (FT) cells. (**B**) Mean action potential durations at 30, 50, 80 and 90 % repolarization (APD_30_, APD_50_, APD_80_ & APD_90_ respectively). (**C**) Mean resting membrane potential (RMP) and action potential amplitude (APA). Bars in graphs indicate standard error of the mean. N indicates number of animals; n, number of cells; and ns, not significant. Unpaired t-test and One Way ANOVA: ns, *p<0.05.

## DISCUSSION

In this study, we present a three-tiered strategy encompassing visual/manual, automatic, and AI-driven methods to identify and classify cells isolated from the LVFW and FTs from adult sheep hearts. Firstly, FT cells were isolated using two successive dissociation methods applied to the central sections of FTs, without connection with the valvular cups, which are known to support primarily the ventricular conduction system^9,10,11^. The first method involved perfusing the coronary system with an enzymatic solution, while the second method required a mechanical chunk approach with an enzyme-free 0.2 mM Ca^2+^ solution. Secondly, FT cells were distinguished from other cardiac cell types using a multifaceted approach, including morphological selection, an AI-based identification model, immunostaining, structural studies, and electrophysiological approach. Our results indicate that expert-guided visual inspection to identify FT cells versus LVFW cells was reinforced using automatic and quantitative phenotyping and, furthermore, a deep learning approach could be trained to robustly classify the two cell populations. Overall, FT cells exhibited distinct properties, including cellular morphology, a unique surface of membrane striations, preferential Tbx5 and Cx40 mRNA expression, and a low organization and density of T-tubules. Moreover, cells were subsequently found to have specific electrical properties consistent with previous studies.

### Anatomy

The PF network, located and distributed in the sub-endocardial region in ventricles, is more developed in the LV than in the RV. The asymmetrical distribution of the PF bundle between both ventricles is essential for optimal ventricular activation and contraction. This appears to be an evolutionary adaptation stemming from the heart’s developmental origins^40^. Indeed, the LV generates higher pressures and has a greater workload, requiring a more extensive PF network for rapid electrical propagation and synchronized contraction according to the ventricular mass ^40^. A recent study in rabbits with congestive heart failure showed significant LV remodeling and dilatation, highlighting the need for a well-developed PF network to ensure coordinated contraction under increased workload ^41^. According to this example, a better understanding of this specialized network at the cellular level is needed to unmask its key role in the physiology of sheep heart. Note that the regional characterization of the electrical properties of FT cells was not part of this investigation and requires further studies.

### Cell morphology, structure and mRNA quantification

We used a visual discrimination approach to selectively identify FT cells based on their distinct morphological features compared to LVM. It is also important to emphasize the robustness of our dissociation method, as each experiment consistently yielded both LVM and FT cells suitable for structural or functional analysis. However, it’s noteworthy that the proportion of viable FT cells was considerably lower compared to LVM, due to the rarity of these cellular populations and overall lower available mass of FTs. FT cells were morphologically identified using established visual criteria from the literature, which closely match the phenotype of PF cells, such as elongated/slender shapes^15,16,17,18^, finger-like projections^19^, curves^20,21,22^, tortuous/tormented^15,22,23^ and including spurs as a novel finding of our study. This approach helped to provide a detailed description of these morphological features and to contextualize standardized terminology for this cell population, ultimately improving FT cell identification and broader application of our methods. Within the FT cell population, we did not observe cells meeting morphological criteria previously reported for transitional PF cells^42^. An additional qualitative observation of note was lower intensity myocardial striation signals for FT cells under the same brightfield imaging conditions, compared to LVMs (data not shown). The presence of striations within the cytoplasm of FT cells indicated sarcomeric organization of myofilaments, suggesting their ability to contract although this is not their primary function^43,44^. It was reported that this structural feature facilitated rapid propagation of electrical impulses throughout the heart. Then, to enhance the discrimination between cell groups, we also conducted a comparative study of cell surfaces using cellular imaging and deep learning-derived cellular segmentation approaches. This was coupled with in vitro patch clamp techniques to evaluate plasma membrane surface area as a function of cellular capacitance (Figures 3, 4 & S5 and table S1). The results indicate that FT cells have significantly smaller width, surface area and membrane capacitance compared to LVM, a finding in line with several previous investigations^13,17^, although some contradictory conclusions were found, due to variations in experimental approaches and species investigated^25,45,46,47,48,49^. The T-tubular organization and density of FT cells were also determined by a di-8-ANEPPS staining. It was observed that FT cells exhibit a markedly lower T-tubule organization and density compared to LVM, consistent with the study realized by Sommer and Johnson in 1968, which involved guinea pig, dog and goat^50^. Interestingly, this specific pattern appears to be consistent across species^47^ and the restricted presence of membrane invaginations may lead to less efficient excitation-contraction coupling and also plays a primary role in rapid conduction electrical impulses^51^. Finally, we confirmed by q-PCR that both Tbx5 and Cx40 have a higher mRNA expression level in FT cells than in ovine LVM, as reported in other recent studies^44,52,53,54^.

This observation is also in line with previous investigations indicating that Tbx5 drives Nav1.5 expression and indirectly regulates the function of the cardiac conduction system, which may contribute to the observed higher maximum conduction velocity and amplitude of AP in FT cells compared to LVM^55^. Indeed, despite the lack of statistical significance, this insight may offer an explanation for the slightly increased conduction velocity and AP amplitude observed in FT cells, (Figures 7 & S4A and B).

### Cardiac-cell phenotype characterization and discrimination using deep learning and image processing

Our study represents an innovative experimental approach to improve the accuracy and efficiency of FT cell identification and classification by integrating an automated AI model from a microscopy image. By training the AI-model on our dataset of FT cell morphological features (annotated images), we have significantly increased the reliability in distinguishing FT cells from LVMs. This deep learning approach holds great promise for cellular discrimination and minimizing the potential experimental bias related to cross-contamination of cells, as well as misinterpretation of morphological features. Traditional methods of PF identification often rely on manual visual inspection, which can be subjective, time-consuming and potentially a major source of errors. In contrast, the AI-based approach will provide an objective, high-throughput means of accurately classifying FT and LVFW cells, which can feasibly be extrapolated in future studies to discriminating cell types dissociated from more heterogeneous tissue samples, including endocardial surface and transmural regions of the PF network. Indeed, the deep learning and mathematical development, and implementation of this structural AI-model represents a significant advancement in the field of cardiac cellular electrophysiology research. By enabling the reliable isolation of FT cells populations, we can now conduct more targeted investigations into the unique structural, functional, and electrophysiological properties of these specialized conduction system. Additionally, improving access to cellular constituents of the PF network holds promising perspectives for a better understanding of the molecular mechanisms underlying cardiac arrhythmias and opens potentially the door to new therapeutic approaches. Considered collectively, the integration of an automated structural AI-model for FT cell identification and classification represents a technological breakthrough in the field of cardiac cellular research. This innovative approach promises to enhance the accuracy, efficiency, and reproducibility of FT cell subgroup identification and beyond, ultimately enabling more comprehensive investigations into the role of these specialized cells in cardiac physiology and pathology.

### Electrophysiology

Our results are consistent with the literature, showing a longer AP and a tendency towards higher maximum conduction velocity and amplitude in FT cells. By comparing the AP profiles of these two cell types, we aim to elucidate the underlying cellular mechanisms that govern their unique electrophysiological signatures. Conversely, although LVM constitutes the majority of the myocardium (∼75 % of the heart mass), compared to electrophysiological properties of the PF network identified in other species, they exhibit different electrical behaviors characterized by shorter AP durations and lower maximal conduction velocities, and amplitude, which are vital for effective contractility and coordinated beating^56^. Indeed, the prolonged AP duration characterized by longer APD_50_ and APD_90_ found in FT cells in our study, compared to LVM, contributes to maintaining a regular heart rhythm by preventing premature reactivation of the PF network during the refractory period^5,18,57^. This unique electrical property is indirectly linked, as well, to the dynamics of transient Ca^2+^ currents in FT cells^58,59,60^, a facet not explored in the present study. Future investigations are needed to elucidate the relationship between the prolonged AP duration and dynamic of transient Ca^2+^ currents within our two experimental groups. Finally, it is noteworthy that we did not observe significant changes in the resting membrane potential of our two experimental groups, which is a confirmation of the accuracy of our data description. This regional comparative analysis holds the potential to significantly advance our understanding of heart function within our animal model and provides valuable insights for cardiac physiology. Then, our functional data, gathered through experimentation and analysis, shows that the initial rapid depolarization phase (phase 1) and the subsequent plateau phase of AP (phase 2) in FT cells displays similar profiles to those observed in LVM (Figure S3). Furthermore, we observed a consistently prolonged action potential duration in FT cells compared to LVMs. Interestingly, this contrasts with prior findings in other species, where PF cells often exhibit a more negative plateau phase than ventricular myocytes^14,61,62^. Such differences highlight potential species-specific electrophysiological traits or regional heterogeneities within the PF network. These observations underscore the need for further investigation into the ionic mechanisms, particularly repolarizing K⁺ and Ca²⁺ currents that may drive these disparities. Notably, the integration of AI-based image classification in our workflow provides a scalable and objective platform for analyzing rare cardiac cell types. This approach enhances the precision and efficiency of morphological, molecular, and functional characterization, ultimately enabling deeper insights into the physiology and pathophysiology of the ventricular conduction system.

### General Clinical Implications

The methods developed in this study have the potential to advance our understanding of the mechanisms underlying cardiac arrhythmias and conduction disorders in humans. By elucidating the unique electrophysiological signatures of cells related to the Purkinje network, our findings will contribute to the future development of targeted therapeutic strategies for the treatment of human cardiac diseases. Efficient and effective classification of critical cell populations in cardiac arrhythmia, notably the Purkinje network, will pave the way to more targeted therapeutic treatments and novel cellular detection technologies for better guiding clinical interventions aimed at restoring normal electrical conduction in humans.

### Study limitations and future directions

While our method for cardiac FT cell dissociation from adult sheep hearts provided valuable insights into cardiac electrophysiology and give us indirectly interesting perspectives to better understand the intricate crosstalk between cardiac ventricular tissues and PF network in physiological conditions, several limitations should be discussed. (1) Although efforts were made to ensure consistency in the dissociation procedures, inherent variability in experimental and biological parameters among individual hearts could have influenced the results. Furthermore, the use of control sheep hearts may not fully recapitulate the complexities of human cardiac physiology and diseases^25,36^. Therefore, further validation of this method on pathological sheep models involving the PF network, or human tissue is needed. (2) Although our algorithm performs well in distinguishing FT cells from LVM, we cannot guarantee the total purity of our cell preparations and cannot exclude the possibility that some LVMs and FT cells share similar morphological characteristics. The exploration of novel specific biomarkers holds potential for enhancing the discrimination and characterization of these rare phenotype cells. Indeed, while Tbx5 and Cx40 mRNA quantification shows promise in distinguishing FT cells, with or without subgroup distinction, integrating these markers with additional biomarkers will not only deepen our understanding but also enhance the quality of our analysis. Nevertheless, further investigations are needed to correlate our observations at the mRNA level with protein expression, cellular shape and functional outcomes. While automated AI-model segmentation of our two experimental groups offers significant advantages in terms of efficiency, reproducibility, and consistency, it is important to keep in mind the limitations of such an approach. Our AI-model learns to distinguish our cell types using a dataset of annotated images derived from morphological analyses. Therefore, it is crucial to improve our dataset by adding new sets of images from morphological analysis and specific staining of new and established biomarkers. This will strengthen the model’s learning and mitigate inappropriate detections. (3) The lack of interaction with the deep learning tool at this stage of development could also limit our level of interpretation. Indeed, it would be pertinent to develop and optimize our program to better interact with the AI (AI explainable) in order to extract morphological features from images used to classify the different cell types and ultimately provides new ones, difficult to identify by the human eye. Additionally, imaging factors, such as image resolution, sensitivity of camera detectors may lead to differences in the algorithm performance and capacity to discriminate cell populations. Further analysis of imaging sensitivity will enhance the overall effectiveness of our analysis. (4) We have not investigated the ionic bases of the AP in order to elucidate the underlying mechanisms of AP generation in both cell types. A detailed characterization of the involved ionic currents could also provide valuable insights into the phenotype, physiological functions and potential pathological alterations.

## CONCLUSION

This study presents a comprehensive and rigorously validated methodology for the isolation, characterization, and classification of PF cells from adult sheep hearts. By combining qualitative visual assessment, conventional image-based morphometrics, and high-precision deep learning via YOLOv9, we establish a robust, multi-layered pipeline capable of accurately distinguishing PF cells from LVMs within heterogeneous cardiac tissues. This integrative approach not only enhances classification accuracy but also improves the consistency and scalability of PF cell identification; an essential advancement given the structural complexity and low abundance of these cells.

Our findings confirm that FT-dissociated cells exhibit distinct morphological, structural, transcriptional, and electrophysiological features consistent with PF cell identity. The successful application of deep learning to automate phenotype recognition represents a key scientific innovation, paving the way for high-throughput, objective analysis of specialized cardiac cell populations. Importantly, this work addresses longstanding technical challenges in PF research and provides a foundation for future investigations into the role of the Purkinje network in arrhythmogenesis and SCD.

Altogether, our methodology enables a more precise exploration of the ventricular conduction system and offers a powerful new toolset to advance cardiac electrophysiology, disease modeling, and potential therapeutic targeting of conduction-related pathologies.

## Acknowledgements

The authors wish to express their gratitude to the animal facilities staff: Virginie Loyer, Gabrielle Carpentier, Stéphane Bloquet, Agathe Vallespir, John Aleksander-Haffreingue, Benjamin Péré, Céline Ayez and Coralie Lecompte.

## Institutional Review Board Statement

This study was carried out in accordance with the UE Directive 2010/63/EU for the protection of animals used for scientific purposes and approved by the local ethical authorities (CEEA50) at the University of Bordeaux, France.

## Informed Consent Statement

Not applicable.

## Data Availability Statement

The data supporting the study findings can upon reasonable request be made available from the corresponding authors

## Author contributions

Conceptualization, S.C., R.W., M.H., and O.B.; methodology, S.C., C.RM., A.EG., R.W., and O.B.; Conceptual design and development of the software, S.C., A.EG. respectively; formal analysis, T.H., K.M., and S.C.; investigation, S.C., T.H., K.M., P.P., F.C., S. Charron., F.B. and A.EG.; resources, M.H., and O.B.; writing-original draft preparation, S.C.; writing-review and editing, T.H., K.M., A.EG., R.G., D.B., M.H., R.W., O.B. supervision R.W., and S.C. All authors have read and agreed to the published version of the manuscript.

## Supplemental Material

Tables S1 & S2

Figures S1–S5

## Nonstandard Abbreviations and Acronyms

AP: Action Potential
CMS: Cardiomyocytes
FTs: False tendons
LV: Left ventricle
LVM: Left Ventricular Myocyte
PBS: Phosphate Buffer Saline
PF: Purkinje Fiber
RV: Right Ventricle
SCD: Sudden Cardiac Death
VF: Ventricular Fibrillation
VT: Ventricular Tachycardia

