## Supplementary material for "AI-based identification of cardiac Purkinje fiber cells isolated from whole adult sheep hearts": Supplemantal data

### Supplemental data

#### Tables Supplemental data

| Cell parameters | LVM | FT cells | $P_{value}$ |
|---|---|---|---|
| **Length (µm)** | 117.4 ± 4.2 | 116.2 ± 4.6 | 0.8 |
| **Width (µm)** | 37.0 ± 2.6 | *22.6 ± 1.2 | <0.0001 |
| **Area ($\mu m^2$)** | 3835.2 ± 121.0 | *2603.2 ± 159.4 | <0.0001 |
| **n** | 69 | 72 | |
| **N** | 5 | 6 | |
| **Capacitance (pF)** | 143.9 ± 5.8 | *116.3 ± 4.6 | <0.0001 |
| **n** | 72 | 93 | |
| **N** | 6 | 16 | |

**Table S1: Cardiac cell dimensions (length & width (µm)), surface area ($\mu m^2$) and capacitance (pF).** A comparative investigation was done according to the length, width, the area and capacitance for left ventricular myocytes (LVM) vs false tendon (FT) cells. Corresponding P values are provided in the right column.

| AP parameters | LVM | FT cells | $P_{value}$ |
|---|---|---|---|
| **RMP (mV)** | -70.8 ± 1.5 | -72.3 ± 0,9 | 0.40 |
| **APA (mV)** | 106.5 ± 3.1 | 112.3 ± 2.4 | 0.14 |
| **dV/dT max** | 212.8 ± 31.7 | 236.9 ± 21,8 | 0.52 |
| **$APD_{10}$** | 45.3 ± 9.6 | 52.9 ± 9,6 | >0.999 |
| **$APD_{20}$** | 100.9 ± 14.9 | 135.1 ± 15.1 | 0.94 |
| **$APD_{30}$** | 163.9 ± 18.2 | 219.9 ± 18.3 | 0.48 |
| **$APD_{40}$** | 226.9 ± 19.1 | 305.1 ± 21,1 | 0.10 |
| **$APD_{50}$** | 268.6 ± 21.2 | 364.4 ± 24.3 | *0.02 |
| **$APD_{60}$** | 295.6 ± 22.5 | 395.6 ± 25.3 | *0.01 |
| **$APD_{70}$** | 313.4 ± 23,3 | 413.9 ± 25.6 | *0.01 |
| **$APD_{80}$** | 325.7 ± 23.9 | 425.8 ± 28.8 | *0.01 |
| **$APD_{90}$** | 336.8 ± 24.7 | 435.7 ± 26.0 | *0.01 |
| **n** | 13 | 17 | |
| **N** | 6 | 6 | |

**Table S2: Action potential parameters:** Comparison of action potential parameters from left ventricular myocytes (LVM) and false tendon cells (FTs). Mean resting membrane potential (RMP), action potential amplitude (APA) and maximal upstroke velocity [dV/dtmax (V/s)], are provided. Finally, mean action potential durations at x % of repolarization (APDx) were also reported. Data were compared between LVM and FT cells using ordinary one-way ANOVA / Sidak's multiple comparisons test according to the normality test: * $p < 0.05$. Corresponding P values are provided in the right column.

**Figure supplemental data:**

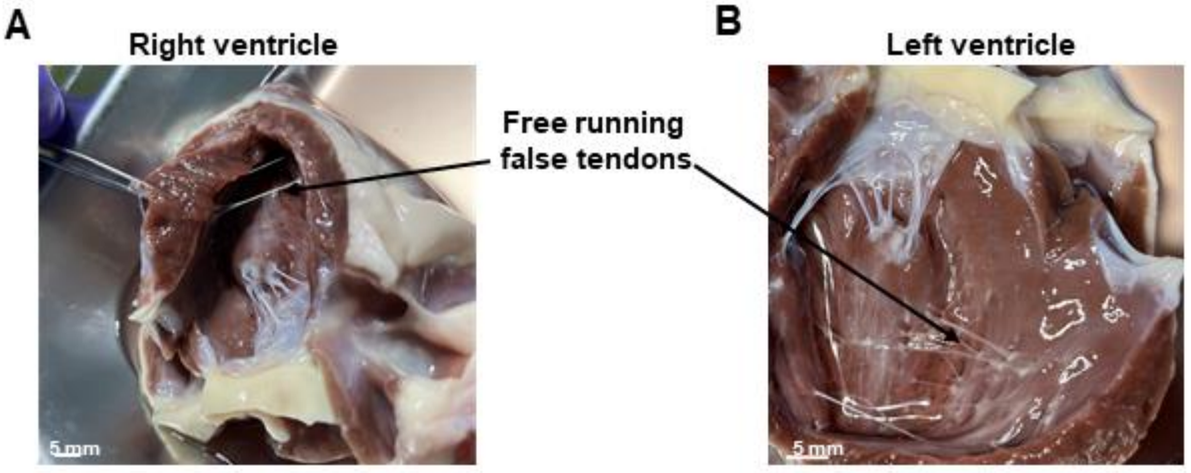


**Figure S1: Photomicrographs of False tendons network on a right (A) and left ventricular (B) strip**. As reported, the left ventricular has a more developed PFs network compared to the right ventricle area.

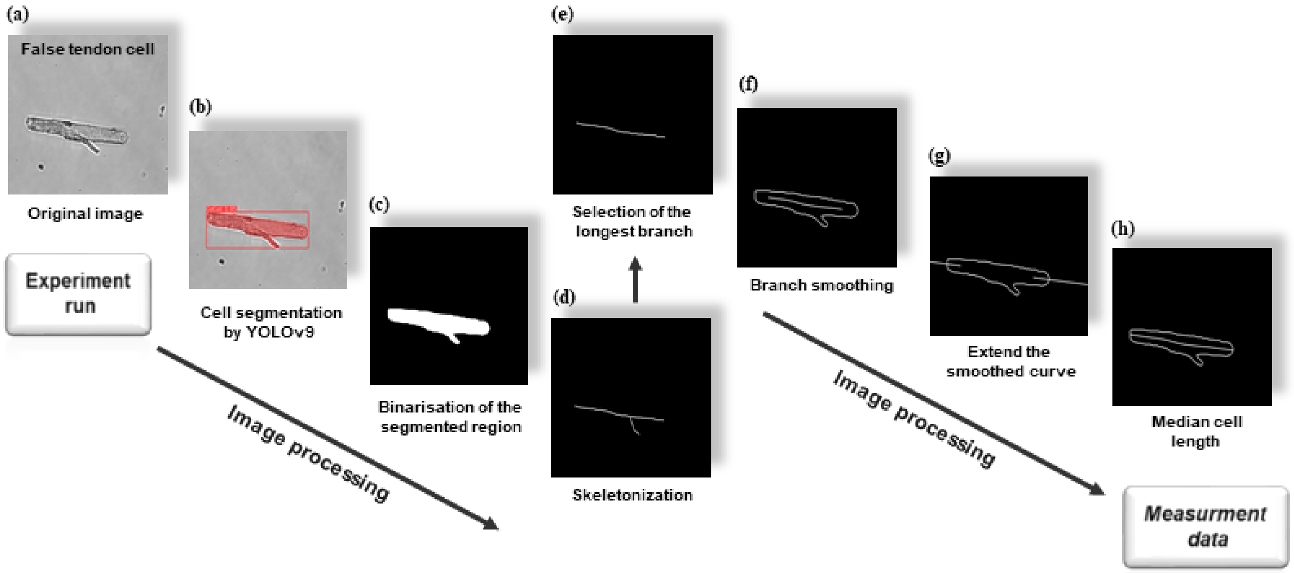


**Figure S2: The procedure for identifying automatically the cell within the image, and determining its area and length:** (a) Original image; (b) Cell detection using YOLOv9; (c) Binarisation of the segmented region; (d) Skeletonization (Lee method); (e) Selection of the longest branch; (f) Branch smoothing; (g) Extend the smoothed curve; (h) Final median cell length curve.

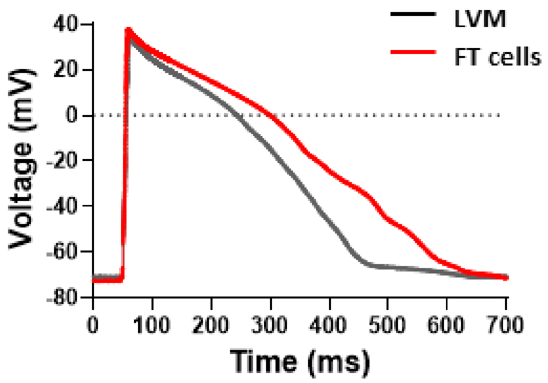


**Figure S3: Average action potential traces for each investigated group.** To assess the presence of a lower plateau in false tendon (FT) cells compared to the left ventricular myocytes (LVM), individual representative AP from the figure 7 were averaged. The results suggest that the plateau of FT cells AP is not lower than LVM AP, under our conditions.

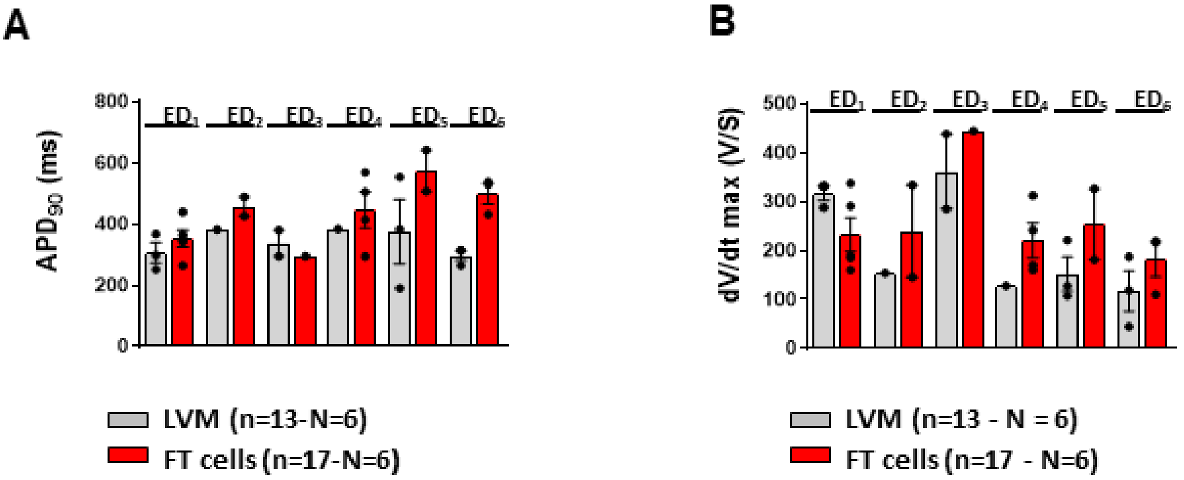

**Figure S4: Distribution of the $APD_{90}$ and dV/dt max across experimental days and cell groups (left ventricular myocytes (LVM) vs. False tendon cells (FT cells)).** Each "$ED_x$" label represents a specific experimental day, with "x" indicating the day number. Data are represented as mean ± standard error of the mean with individual values for each cell. N indicates number of animals; n, number of cells.

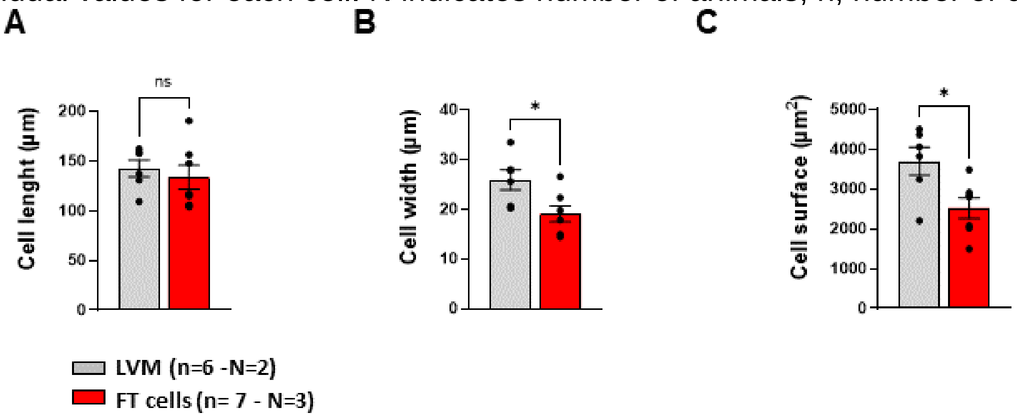


**Figure S5: Cell parameters from left ventricular myocytes (LVM – grey) and False tendon cells (FT cells – red) in sheep.** AI analysis was employed to classify cell phenotypes using a series of images captured during the functional recordings (cf. Figure 7). **(A)** cell length (µm), **(B)** cell width (µm), and **(C)** cell surface (µm²). Data are represented as mean ± standard error of the mean with individual values for each cell. N indicates number of animals; n, number of cells; and ns, not significant. Unpaired t-test: ns, *p<0.01.
